# Subclinical Atherosclerosis is Associated with Discrepancies in BAFF and APRIL Levels and Altered Breg Potential of Precursor-like Marginal Zone B-Cells in HIV Treated Individuals

**DOI:** 10.1101/2022.08.26.505170

**Authors:** Matheus Aranguren, Kim Doyon-Laliberté, Mohamed El-Far, Carl Chartrand-Lefebvre, Jean-Pierre Routy, Jean Guy Barril, Benoît Trottier, Cécile Tremblay, Madeleine Durand, Johanne Poudrier, Michel Roger, Canadian HIV and Aging Cohort Study

## Abstract

Chronic inflammation persists in people living with HIV (PLHIV) despite antiretrovial therapy (ART), and is involved in their premature development of cardiovascular diseases (CVD) such as atherosclerosis. We have previously reported that an excess of “B-cell activating factor” (BAFF), an important molecule for the selection and activation of first line Marginal Zone (MZ) B-cell populations, is associated with deregulations of precursor-like MZ (MZp), whose potent B-cell regulatory (Breg) capacities are altered in PLHIV, early on and despite 1-2 years of ART. Based on these observations, and growing evidence that MZ populations are involved in atherosclerosis control, we designed a cross sectional study to explore the associations between BAFF and its analogue “A proliferation-inducing ligand” (APRIL) with subclinical CVD in long time treated individuals of the Canadian HIV and Aging Cohort Study (CHACS) imaging sub-study group. We also characterized the Breg profile of MZp from the blood of these individuals. Results were correlated with the total volume of atherosclerotic plaques (TPV) and with CVD risk factors and biomarkers. TPV was measured using cardiac computerised tomography angiography, and presence of CVD was defined as TPV > 0. We report that blood levels of BAFF are elevated and correlate positively with CVD and its risk factors in PLHIV from the CHACS, in contrast to APRIL levels, which correlate negatively with these factors. Expression levels of Breg markers such as NR4A3, CD39, CD73 and CD83 are significantly lower in PLHIV when compared to those of HIV-uninfected controls. *In vitro* experiments show that APRIL upregulates the expression of Breg markers by blood MZp from HIV-uninfected individuals, while this modulation is dampened by the addition of recombinant BAFF. Altogether, our observations suggest that strategies viewed to modulate levels of BAFF and/or APRIL could eventually represent a potential treatment target for CVD in PLHIV.

## Introduction

Antiretroviral therapy (ART) has been a staple in the treatment of Human Immunodeficiency Virus (HIV) infection since its appearance in the early 90s. It is considered one of the revolutions of the 20^th^ century in medicine, effectively converting a deadly disease into a manageable condition, and giving people living with HIV (PLHIV) a quasi-normal life expectancy [1]. In 2015, it was estimated that 50% of all PLHIV in Canada were over 50 years old and were thus more commonly being afflicted by aging-associated diseases, some of which are developing prematurely (5 to 10 years earlier than the general population), including cardiovascular diseases (CVD) [2, 3]. Indeed, it has been shown that PLHIV are more prone to developing atherosclerosis plaques, the primary cause of heart disease, when compared to healthy individuals, even after adjustment for traditional CVD risk factors such as smoking, hypertension and diabetes [4, 5]. The root cause of this development is not well understood, but ART toxicity, chronic inflammation and pathogenic opportunistic infections have been suggested to contribute to premature CVD [2, 3, 6]. Given the importance of inflammation in the formation of atherosclerosis plaques, chronic inflammation in the HIV context is therefore considered to be a powerful trigger for this process [7, 8].

Despite the success of ART, chronic inflammation associated with HIV infection persists due to factors such as viral transcription by the latent reservoir, microbial translocation, and long-term immune deregulation [9-11]. We have shown that B-cell activating factor (BAFF) is in excess in the blood of PLHIV from the Montreal Primary HIV Infection Cohort (PHI) and correlates with viral factors such as Nef, and is concomitant with inflammatory markers such as elements of microbial translocation [12]. Moreover, this excess is detected as early in the acute phase and persists despite 1-2 years ART [13] (Doyon-Laliberté, Aranguren *et al, unpublished observations*). BAFF is a critical B-cell survival and differentiation factor and helps shape the innate marginal zone (MZ) B-cell pool by contributing to MZ selection during normal B-cell ontogeny [14, 15]. The excess of BAFF we described in PLHIV from the Montreal PHI cohort was concomitant with hyperglobulinemia and deregulations of the B-cell compartment, notably with increased frequencies of innate MZ precursor-like (MZp) B-cells [13]. We reported similar observations for HIV-infected Beninese female commercial sex workers, SIV-infected macaques and initially for HIV-transgenic mice [16-18], suggesting that excess BAFF and deregulated MZ populations are reliable markers of inflammation in the context of HIV.

BAFF is mainly produced by neutrophils, myeloid cells such as monocytes and dendritic cells, activated T-cells and B-cells, and the stromal cells of secondary lymphoid organs. BAFF can be found in a soluble form, or in a membrane-bound form. A Proliferation-inducing Ligand (APRIL) is a BAFF analogue, which can only be found in a soluble form or complexed with heparin sulfate proteoglycans (HSPG). BAFF can signal through three receptors, BAFF-Receptor (BAFF-R), Transmembrane Activator and CAML Interactor (TACI), and B-cell Maturation Antigen (BCMA), while APRIL only signals through the latter two receptors [15]. Although they share similar roles, BAFF and APRIL also have different functions in B-cell immunity, as observed in their individual knock-out phenotypes [19, 20].

We have previously shown that in HIV-uninfected individuals, MZp present a powerful B-cell regulatory (Breg) potential and function [21, 22]. Indeed, *ex vivo*, MZp strongly express IL-10 as well as the transcription factors Nuclear Receptors (NR)4A1, NR4A2 and NR4A3, which are heavily involved in anti-inflammatory responses. MZp also express the immunoregulatory molecule CD83 [23], whose expression is dependent on NR4As and whose signalling we have shown to be important for MZp Breg function [21]. Moreover, MZp also express the ectonucleotidases CD39 and CD73, involved in the conversion of the pro-inflammatory ATP into the anti-inflammatory adenosine (ADO) [21, 24]. Importantly, recent observations show that blood MZp Breg capacities are highly deregulated in PLHIV from the Montreal PHI cohort, involving downregulation of the immunoregulatory markers NR4A1-3, CD83 and CD73 as well as immune checkpoint molecule Programmed Death Ligand 1 (PD-L1), likely contributing to their altered Breg function. Of note, this deregulation was not restored by ART. Furthermore, MZp loss of Breg capacity appears to be directly related to the excess of BAFF, as *in vitro*, excess BAFF downregulates the expression of NR4A1 and NR4A3, as well as CD83, and alters MZp Breg function (Doyon-Laliberté; Aranguren *et al, unpublished*).

The role of B-cells in atherosclerosis formation is complex and subset-dependant and has been mostly studied in murine models [25]. On one hand, follicular (FO) B-cells are considered atherogenic by generating germinal center (GC) responses and IgG directed against oxidised low density lipoprotein (oxLDL), and this process correlates with CVD [26]. Moreover, FO B-cells are also involved in the formation of tertiary lymphoid organs at the *tunica adventitia* of blood vessels, also affected by atherosclerosis, allowing for the generation of GC and production of anti-oxLDL IgG *in situ* [26-28]. On the other hand, MZ B-cells are considered atheroprotective as they mitigate follicular helper T-cell function in GC via PD-1/PD-L1 interactions [29]. Interestingly, the atheroprotective role of MZ is dependent on the expression of NR4A1, as its deletion in mice aggravates atherosclerosis development due to the downregulation of PD-L1 [30]. Moreover, anti-oxLDL IgM, as opposite to IgG, is considered atheroprotective. As such, the fact that MZ B-cells are massive producers of IgM [14, 26] could be related to their atheroprotective role.

Based on these observations, our aim was thus to study the association between BAFF and APRIL levels, as well as MZp deregulation, with the presence of early subclinical CVD in PLHIV from the Canadian HIV and Aging Cohort Study (CHACS). We show that BAFF levels remain relatively elevated in the blood of PLHIV from the CHACS despite 15 or more years of ART, while APRIL levels are not affected. Furthermore, BAFF levels correlate positively with CVD risk factors, while APRIL levels correlate negatively with these factors. Moreover, BAFF and APRIL levels correlate negatively with each other. We also show that the Breg profile of MZp from the blood of long term treated PLHIV from the CHACS are deregulated, as expression levels of NR4A3, CD39, CD73 and CD83 are significantly lower when compared to those of HIV-uninfected controls. In accordance with a seemingly atheroprotective role, *in vitro* experiments show that recombinant APRIL upregulates the expression of Breg markers such as NR4A1, NR4A3 and IL-10 by blood MZp from HIV-uninfected individuals, while this modulation is dampened by the addition of recombinant BAFF. Altogether, our results shed light on a possible atheroprotective role of APRIL in shaping MZp Breg capacities, in contrast to that of excess BAFF, likely more atherogenic and associated with altered Breg capacities.

## Materials and Methods

### Specimens and clinical data collection

We herein conducted a cross sectional study, using data and samples from the CHACS imaging sub-study group. Briefly, the CHACS recruits PLHIV having either lived with HIV for at least 15 years or are over the age of 40, as well as HIV-uninfected controls over the age of 40. Controls are recruited at the same clinics as the PLHIV participants, and the full study protocol and data collection forms have been published [6]. Participants from the CHACS cohort who were free of overt cardiovascular disease (had never suffered a myocardial infarction, coronary revascularization, angina, stroke of peripheral vascular revascularization) and had a 10-year Framingham risk score (the probability of developing CVD in the near future) ranging from 5 to 20% were invited to participate in the cardiovascular imaging substudy [31]. Data on all traditional cardiovascular risk factors was collected prospectively as part of the CHACS visits.

For the present study, we have randomly selected 79 participants from the CHACS cardiovascular imaging prospective sub-study, which included successfully ART-treated PLHIV with or without subclinical CVD and HIV-uninfected controls with or without subclinical CVD. Presence/absence of subclinical CVD was defined as the presence or absence of coronary artery atherosclerosis plaques, which were measured as described below. Four study groups were generated as follows: 20 HIV uninfected participants without CVD (HIV-CVD-), 20 HIV uninfected participants with CVD (HIV-CVD+), 20 HIV infected participants without CVD (HIV+ CVD-), and 19 HIV infected participants with CVD (HIV+ CVD+).

### Characterisation of atherosclerosis plaques

Coronary plaques were measured as previously described [31, 32]. Briefly, individuals recruited from the CHACS were administered 50-75 mg metoprolol (a beta blocker used to lower heart rate) orally when heart rate was higher than 60 beats per minute. Then, coronary computerized tomography (CT) angiography was performed using a 256-section CT scanner (Brilliance iCT; Philips Healthcare) and 370 mg/ml iopamidol as a contrast (Bracco Imaging) at a rate of 5 ml/sec after bolus tracking. Electrocardiogram-gating was used. Slice thickness was 0.625 mm. Plaque images were assessed on a patient-by-patient basis by a board-certified radiologist (C.C.L) and measured afterwards using a semi-automated software application (Aquarius iNtuition 4.4.6; TeraRecon). Image assessors were blinded to CVD risk factors and HIV status.

### Measure of soluble BAFF, APRIL and CD83 blood levels

Soluble BAFF, APRIL and CD83 levels were measured in the serum/plasma of participants of the four study groups by using the Human BAFF/BLyS/TNFSF13B Quantikine ELISA Kit (R&D systems), APRIL Human ELISA Kit (ThermoFisher) and Human CD83 DuoSet ELISA Kit (R&D systems), respectively, by following the manufacturer’s protocol. For measuring soluble CD83, samples were diluted 1:3 prior to testing. Concentration of the analytes was determined using Microsoft Excel.

### Flow cytometry characterisation of Breg markers and membrane BAFF expression levels

Cell expression levels of the Breg markers NR4A1, NR4A3, CD83, CD39 and CD73 were assessed by multicolour flow-cytometry on blood samples from the participants of the four study groups. Briefly, 10^7^ Peripheral Blood Mononuclear Cells (PBMCs), which had been cryopreserved until use, were thawed, washed with Iscove’s Modified Dulbecco’s Medium (IMDM, Thermo Fisher) followed by 1x Phosphate Buffered Saline (PBS) pH 7.4 (ThermoFisher) and processed for flow-cytometry. Live/dead exclusion was performed using Aqua-LIVE/DEAD Fixable Stain (Invitrogen Life technologies, Eugene, OR, USA). Non-specific binding sites were blocked using FACS buffer (1x PBS, 2% heat-inactivated FBS (hi-FBS), and 0.1% sodium azide) supplemented with 20% hi-FBS, 10 μg mouse IgG (Sigma-Aldrich, St-Louis, MO, USA) and 5 μg Human BD FCBlock (BD Biosciences). The following conjugated mouse anti-human monoclonal antibodies (mAbs) were used to detect extracellular markers: APC Anti-CD19, BB515 Anti-IgM, BV421 Anti-CD10, BUV395 Anti-CD73, BV786 Anti-CD39, PE-Cy7 Anti-CD83 (BD Biosciences), PerCP-eFluor 710 Anti-CD1c (eBioscience, San Diego, CA, USA) and PE Anti-BAFF (Invitrogen). Intranuclear labelling was performed using the FoxP3/Transcription Factor Staining Buffer Set (eBioscience). Non-specific binding sites were blocked using 20% hi-FBS. The PE-conjugated human REA clone anti-mouse NR4A1 was used and compared to that of PE-conjugated human REA isotype control, as previously described [21]. The PE-conjugated mouse anti-human NR4A3 mAb was from Santa Cruz Biotechnology. Intra-cellular labelling for IL-10 detection was performed using the Intracellular Fixation & Permeabilization Buffer Set (eBioscience), with the PE-mouse Anti-Human IL-10 Ab. Cells were suspended and kept at 4°C in 1.25% paraformaldehyde prior to analysis. Data acquisition was performed with LSRIIB (BD), and analysis was done with FlowJo 10.7.1 software and GraphPad Prism. Gating strategy was based on Full Minus One (FMO) values and isotype controls (Supplementary Figure 1). Anti-mouse Ig(κ) Compbeads and CS&T Beads were used to optimize fluorescence compensation settings and calibrate the LSRIIB, respectively.

### Culture of blood human B-cells with recombinant APRIL and BAFF

Cryopreserved PBMCs from three HIV-uninfected donors were thawed and washed in IMDM. Total blood B-cells were then negatively enriched >95% by an immunomagnetic based technology (Dynabeads Untouched Invitrogen Life technologies). Total B-cells were subsequently cultured at a concentration of 10^6^ cells/ml in IMDM supplemented with 10^−4^ β-2-mercaptoethanol, 10% hi-FBS and 1% penicillin/streptomycin (ThermoFisher) in absence or presence of APRIL (Recombinant Human APRIL/TNFSF13 Protein, R&D systems) at 50 ng/ml, 250 ng/ml or 500 ng/ml, and/or BAFF (Recombinant Human BAFF/BLyS/TNFSF13B Protein, R&D systems) at the same concentrations for 18 hours at 37°C and 5% CO_2_. Brefeldin A was added for the last 4 hours of incubation prior to staining for intra-cellular IL-10. Cells were then recovered and processed for flow-cytometry as stated above.

### Statistical analyses

Statistical significance of differences between groups was assessed with a one-way ANOVA with post-hoc Tukey test for data normally distributed or otherwise with Kruskal-Wallis test with post-hoc Dunn tests for multiple comparisons. For the correlations, a Pearson correlation was used if data was found to be normally distributed, otherwise a Spearman correlation was used instead. For the categorical data, a Fisher’s exact test was used. Analyses were performed using GraphPad Prism 9.1.2, on Windows. Results were considered significant when p < 0.05.

### Ethics Statement

Written informed consent was obtained from all subjects who participated in the study. The study was conducted in accordance with the Declaration of Helsinki. The methods reported in this manuscript were performed in accordance with the relevant guidelines and regulations and all experimental protocols were approved by the Centre Hospitalier de l’Université de Montréal (CHUM) Research Ethics Committees and all other Research Ethic Committee of the CHACS cohort (# CE 11.063).

## Results

### Socio-demographic characteristics of the study cohort

Socio-demographic characteristics of our study cohort can be found in Table 1. Upon characterizing the cohort, we found significant differences for age between the HIV-CVD- (mean of 54.64) and HIV-CVD+ (mean of 61.93) groups (p= 0.015), and between the HIV-CVD+ and HIV+ CVD- (mean of 57.08) groups (p= 0.016), as assessed by one-way ANOVA with post-hoc Tukey. No differences were found between HIV-CVD+ and HIV+ CVD+. Significant differences were found for the LDL levels between the HIV-CVD- (mean of 3.40 nmol/L) and HIV+ CVD- (mean of 2.45 nmol/L) groups (p = 0.0078), and between the HIV-CVD+ (mean of 3.31 nmol/L) and HIV+ CVD-groups (p = 0.026), as assessed by Kruskal-Wallis test with post-hoc Dunn, with no difference between the HIV+ CVD- and HIV+ CVD+ groups. A significant difference was found for the HDL levels between the HIV-CVD- (mean of 1.49 nmol/L) and HIV+ CVD- (mean of 1.17 nmol/L) groups (p = 0.039), as assessed by Kruskal-Wallis test with post-hoc Dunn. No differences were found between the HIV+ CVD- and HIV+ CVD+ groups. Importantly, statin usage is not controlled in the CHACS, and its usage seems to be augmented in PLHIV [31]. However, we did not see any significant difference for statin usage among the study groups.

**Table 1.**
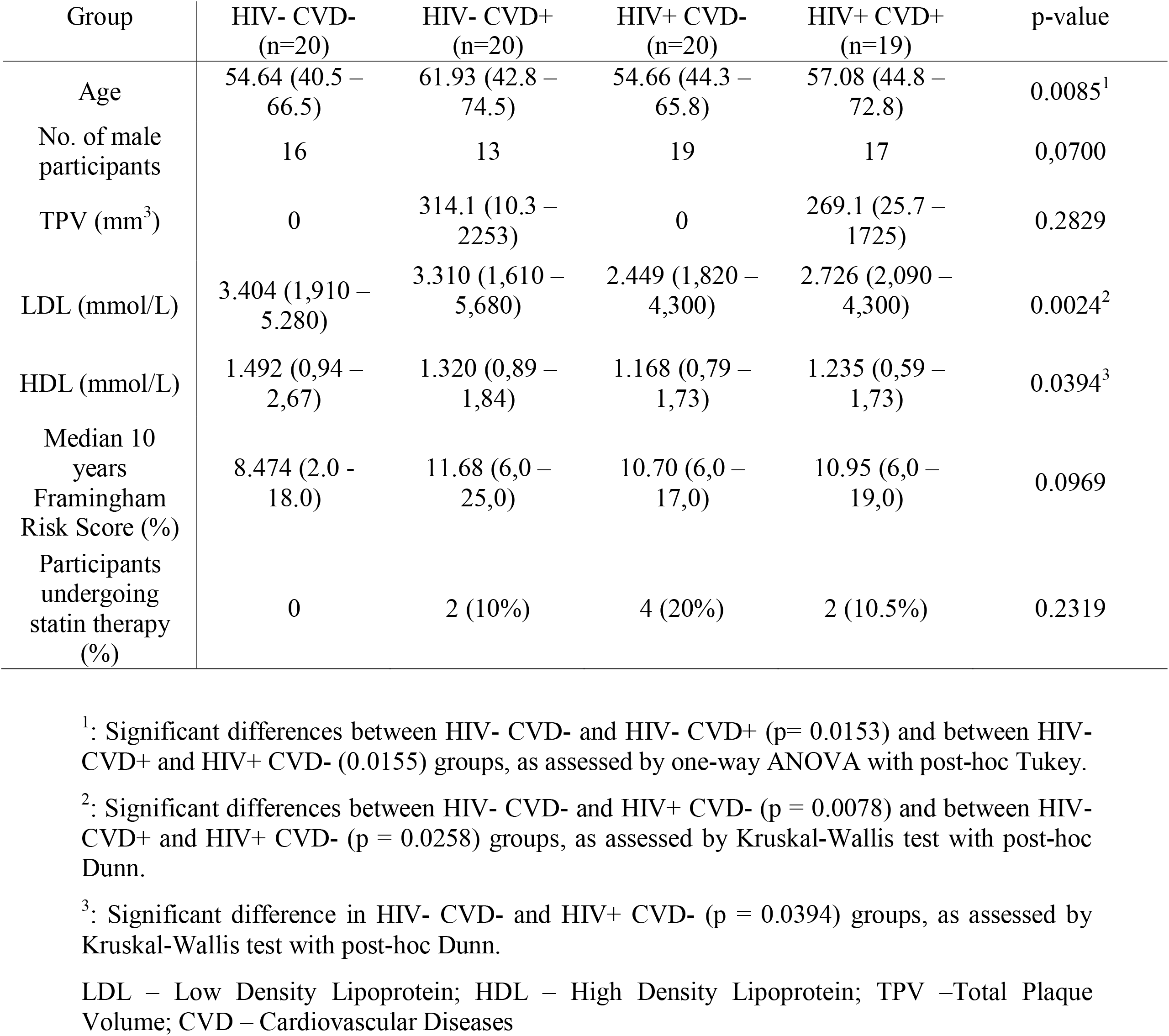
Sociodemographic characteristics of the sub-cohort (A) used in this study.

| Group | HIV- CVD-<br>(n=20) | HIV- CVD+<br>(n=20) | HIV+ CVD-<br>(n=20) | HIV+ CVD+<br>(n=19) | p-value |
| --- | --- | --- | --- | --- | --- |
| Age | 54.64 (40.5 – 66.5) | 61.93 (42.8 – 74.5) | 54.66 (44.3 – 65.8) | 57.08 (44.8 – 72.8) | 0.0085<sup>1</sup> |
| No. of male participants | 16 | 13 | 19 | 17 | 0,0700 |
| TPV (mm<sup>3</sup>) | 0 | 314.1 (10.3 – 2253) | 0 | 269.1 (25.7 – 1725) | 0.2829 |
| LDL (mmol/L) | 3.404 (1,910 – 5,280) | 3.310 (1,610 – 5,680) | 2.449 (1,820 – 4,300) | 2.726 (2,090 – 4,300) | 0.0024<sup>2</sup> |
| HDL (mmol/L) | 1.492 (0,94 – 2,67) | 1.320 (0,89 – 1,84) | 1.168 (0,79 – 1,73) | 1.235 (0,59 – 1,73) | 0.0394<sup>3</sup> |
| Median 10 years Framingham Risk Score (%) | 8.474 (2.0 - 18.0) | 11.68 (6,0 – 25,0) | 10.70 (6,0 – 17,0) | 10.95 (6,0 – 19,0) | 0.0969 |
| Participants undergoing statin therapy (%) | 0 | 2 (10%) | 4 (20%) | 2 (10.5%) | 0.2319 |

<sup>1</sup>: Significant differences between HIV- CVD- and HIV- CVD+ (p= 0.0153) and between HIV- CVD+ and HIV+ CVD- (0.0155) groups, as assessed by one-way ANOVA with post-hoc Tukey.
<sup>2</sup>: Significant differences between HIV- CVD- and HIV+ CVD- (p = 0.0078) and between HIV- CVD+ and HIV+ CVD- (p = 0.0258) groups, as assessed by Kruskal-Wallis test with post-hoc Dunn.
<sup>3</sup>: Significant difference in HIV- CVD- and HIV+ CVD- (p = 0.0394) groups, as assessed by Kruskal-Wallis test with post-hoc Dunn.
LDL – Low Density Lipoprotein; HDL – High Density Lipoprotein; TPV –Total Plaque Volume; CVD – Cardiovascular Diseases

### Excess BAFF levels in the blood of PLHIV from the CHACS correlates with atherosclerosis risk factors

In order to assess the association between BAFF levels and presence of subclinical CVD in PLHIV from the CHACS and HIV-uninfected controls, blood samples (both plasma and serum as well as PBMCs) from the four study groups were screened and compared for their expression levels of soluble and total membrane BAFF. Our data indicate that both HIV+ groups possess higher levels of soluble and membrane-bound BAFF when compared to both HIV-groups. Moreover, we find that the HIV+ CVD+ group presents the highest levels amongst the four study groups (Fig. 1.A, B). We also find a positive correlation trend between total plaque volume (TPV) and blood soluble BAFF levels in the HIV+ CVD+ group (Fig.1.C), and to a lesser extent in the HIV-CVD+ group (Fig. 1.D). Importantly, even though HIV-CVD+ individuals are significantly older than the other groups, we do not see a correlation between BAFF levels and age (data not shown), suggesting that the differences we see for BAFF levels in both HIV-uninfected groups are not due to age differences.

**Figure 1:**
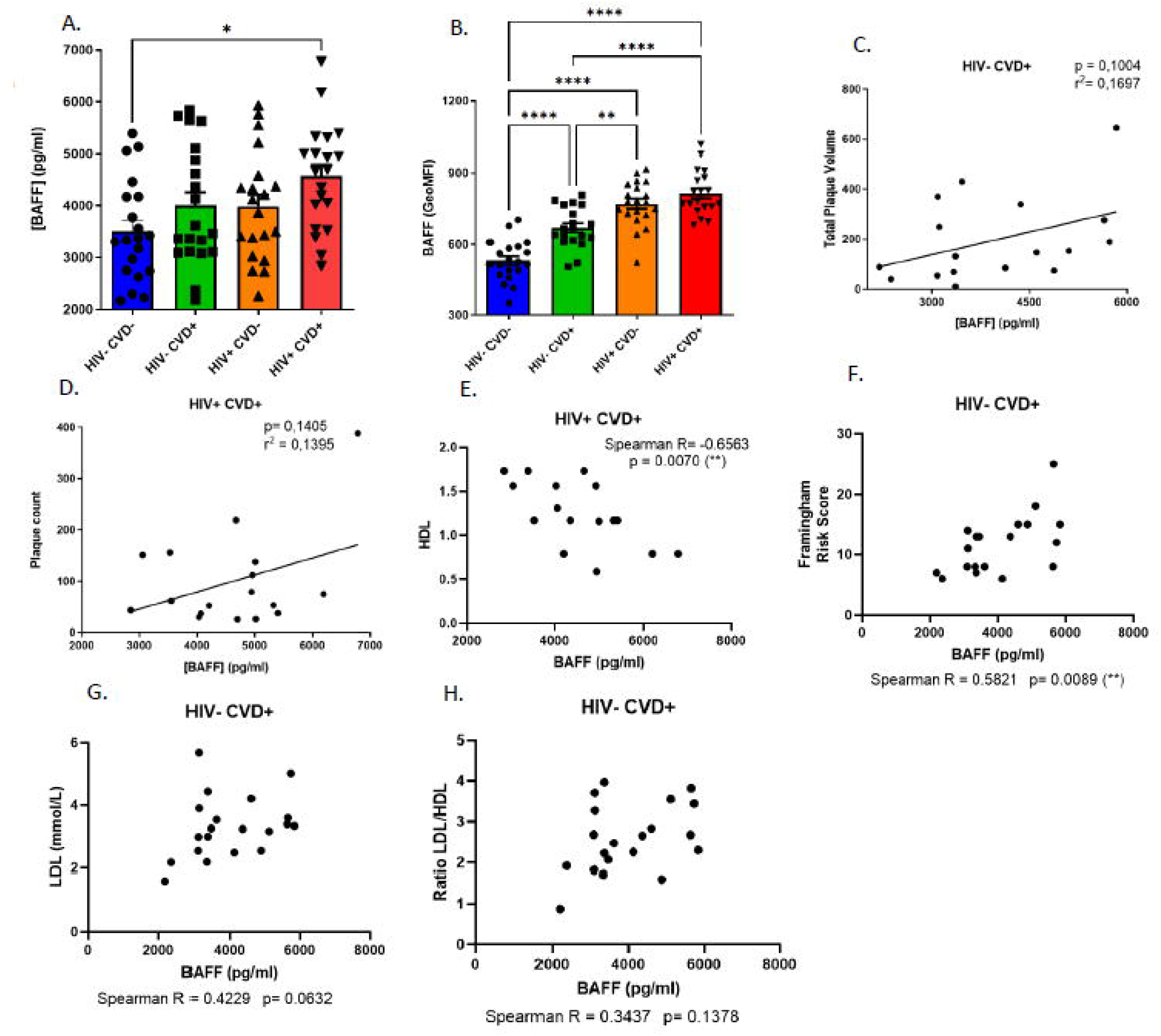
BAFF levels and correlations between soluble BAFF and CVD risk factors in the blood of HIV-uninfected and HIV-infected individuals of the CHACS cohort. Levels of soluble (A) and membrane-bound (B) BAFF in the blood of HIV uninfected participants, without and with CVD (HIV-CVD-, HIV-CVD+, respectively), and HIV-infected participants, without and with CVD (HIV+CVD-, HIV+CVD+, respectively) selected in sub-cohort (A). Correlation between total plaque volume (TPV) and soluble BAFF blood levels in HIV-CVD+ (C) and HIV+CVD+ (D) participants. Correlation between soluble BAFF and HDL (E) blood levels in HIV+CVD+ participants. Correlation between soluble BAFF blood levels and the Framingham risk score (F), LDL (G) and LDL/HDL (H) ratio in HIV-CVD+ participants. Normality was assessed with the Shapiro-Wilk test. A One-Way ANOVA with a post-hoc Tukey was used for testing statistical differences between groups in Fig. 1A, B. A Pearson correlation was used for testing for the correlations in Fig 1C, D. A Spearman correlation was used for testing for the correlations in Fig. 1E, F, G and H. LDL – Low Density Lipoprotein; HDL – High Density Lipoprotein; GeoMFI – Geometric Mean Fluorescence Intensity; CVD – Cardiovascular Disease. * p < 0,05; ** p< 0,01; *** p< 0,001; **** p < 0,0001

We also took advantage of an independent ongoing study of the CHACS imaging sub-study lead by another team (M. E. F) and which will be referred to herein as sub-cohort (B), in contrast to the main cohort described above, referred to as sub-cohort (A). To this end, participants who were also participants of sub-cohort (A) have been excluded. For sub-cohort (B), the four study groups described above were generated with 10 HIV-CVD-, 20 HIV-CVD+, 18 HIV+ CVD- and 39 HIV+ CVD+ (n=87). Socio-demographic data of this sub-cohort (B) can be found in Supplemental Table 1. For the four study groups of sub-cohort (B), soluble BAFF levels were measured in the plasma and quantified with the ultrasensitive Meso Scale Discovery® multiplex kits. The assay was performed according to the manufacturer’s protocol. Data analyses were done with the DISCOVERY WORKBENCH 4.0 Software using curve fitting with 4PL, as previously published [33]. Due to the nature of both techniques and the range of detectable BAFF levels, we decided to keep both sub-cohorts separated instead of pooling them together. As such, and similarly to that observed in sub-cohort (A), we find that in the larger sub-cohort (B), both HIV+ groups present higher levels of soluble BAFF in their blood when compared to both HIV-groups, and the HIV+ CVD+ group presents the highest levels amongst the four study groups (Supp. Fig. 2 A). Furthermore, a positive correlation between TPV and soluble blood BAFF levels in the HIV+ CVD+ group of sub-cohort (B) is also observed (Supp. Fig. 2 B), and to a lesser extent in the HIV-CVD+ group of this sub-cohort (B) (Supp. Fig. 2 C).

Interestingly, in the HIV+CVD+ group of sub-cohort (A), soluble BAFF levels seem to correlate negatively with biomarkers generally considered as atheroprotective, such as HDL levels (Fig. 1.E). In this same sub-cohort, we notice that in the HIV-CVD+ group, soluble BAFF levels correlate positively with the 10 years mean Framingham score risk percentage, LDL levels and LDL/HDL ratio (Figs. 1.F-H). However, soluble BAFF levels of the HIV+ CVD+ group do not correlate with the Framingham score (data not shown). Overall, our data show that BAFF levels are not restored despite long term ART and correlate with the presence of subclinical coronary atherosclerosis in both HIV-infected and uninfected individuals, albeit to a lesser extent in the latter.

### APRIL levels correlate negatively with BAFF and atherosclerosis biomarkers in the blood of PLHIV from the CHACS

Next, we investigated the association of APRIL with the presence of subclinical coronary atherosclerosis in the blood of PLHIV from the CHACS. Surprisingly, we find similar blood levels of APRIL between the four study groups of sub-cohort (A) (Fig. 2.A), which contrasts with the relatively higher APRIL levels that we had previously reported for PLHIV from the Montreal PHI cohort [13]. We find that APRIL, in contrast to BAFF, does not correlate with TPV (data not shown). Differently from BAFF, levels of APRIL correlate positively with atheroprotective biomarkers such as HDL levels (Fig. 2.B) in the HIV+CVD+ group, while they seem to correlate negatively with atherogenic risk factors and biomarkers such as the Framinghan Risk Score in both groups with CVD (Figs. 2.C and 2.D), and with the LDL/HDL ratio (Figs. 2.E) in the HIV+ CVD+ group. Interestingly, blood levels of APRIL seem to correlate negatively with those of soluble BAFF in both groups with CVD, more so in the HIV+ CVD+ group (Figs. 2.F and 2.G). Accordingly, we find that a higher BAFF/APRIL ratio correlates negatively with atheroprotective markers like HDL levels in the HIV+ CVD+ group (Fig. 2.H), while correlating positively with atherogenic markers such as the Framinghan Risk Score and LDL/HDL ratio in this group (Fig. 2.I-J) and even in HIV-individuals (Fig. 2 K-M). Overall, our data show that BAFF outweighs a possible atheroprotective impact of APRIL on the presence of subclinical coronary atherosclerosis.

**Figure 2:**
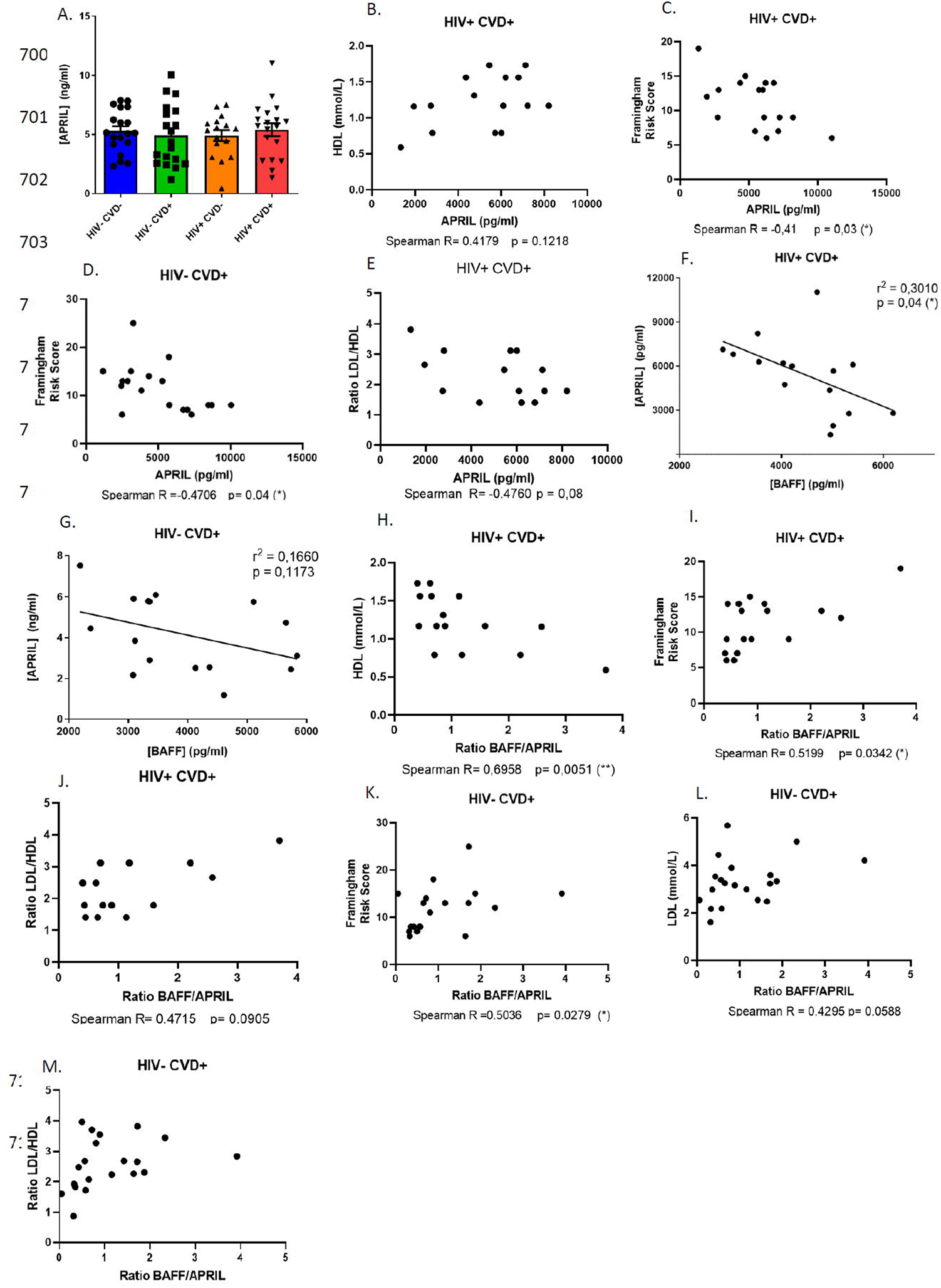
Soluble APRIL levels and correlations between APRIL, BAFF/APRIL ratio and CVD risk factors in the blood of HIV-uninfected (HIV-) and HIV-infected (HIV+) individuals of the CHACS cohort. Levels of soluble APRIL (A) in the blood of HIV uninfected participants, without and with CVD (HIV-CVD-, HIV-CVD+, respectively), and HIV-infected participants, without and with CVD (HIV+CVD-, HIV+CVD+, respectively) selected in sub-cohort (A). Correlation between APRIL and HDL levels (B), Framingham risk score (C) and LDL/HDL ratio (E) in HIV+ CVD+ participants. Correlation between APRIL and Framingham risk score (D) in HIV-CVD+ participants. Correlation between BAFF and APRIL in HIV+ CVD+ (F) and HIV-CVD+ (G) participants. Correlation between BAFF/APRIL ratio and HDL (H), Framingham risk score (I) and LDL/HDL ratio (J) in HIV+ CVD+ participants. Correlation between BAFF/APRIL ratio and Framingham risk score (K), LDL (L) and LDL/HDL ratio (M). Normality was assessed with the Shapiro-Wilk test. A One-Way ANOVA with a post-hoc Tukey was used for testing statistical differences between groups in Fig. 2A. A Pearson correlation was used for testing for the correlations in Fig 2F, G. A Spearman correlation was used for testing for the correlations in Fig. 2B, C, D, E, H, I, J, K, L, M. LDL – Low Density Lipoprotein; HDL – High Density Lipoprotein. CVD – Cardiovascular Disease. * p < 0,05; ** p< 0,01; *** p< 0,001; **** p < 0,0001

### Altered blood MZp Breg capacities in PLHIV from the CHACS

Given that BAFF is found in excess in the blood of long term treated PLHIV of the CHACS, we sought to analyse the impact that this may have on the Breg profile of MZp, as our recent observations suggest that the Breg attributes of blood MZp from PLHIV of the Montreal PHI cohort are altered, despite 2 years of ART (Doyon-Laliberté; Aranguren *et al. unpublished*). In agreement with these observations, upon analysing expression levels of Breg markers by blood MZp of PLHIV from the CHACS and HIV-uninfected controls, we find that while NR4A1 expression levels seem to be similar between the four study groups of sub-cohort (A) (Figs. 3.A and 3.B), those of NR4A3 (Fig. 3.C) and CD83 (Figs 3.D and 3.E) are lower in both HIV+ groups when compared to HIV-groups. Importantly, this downregulation does not seem to vary depending on the CVD status. We also observe an increase in the relative frequencies of MZp in the blood of both HIV+ groups when compared to that observed in both HIV-groups, irrespective of the CVD status (Fig. 3.F), similar to what we have previously reported for PLHIV of the Montreal PHI cohort [22]

**Figure 3:**
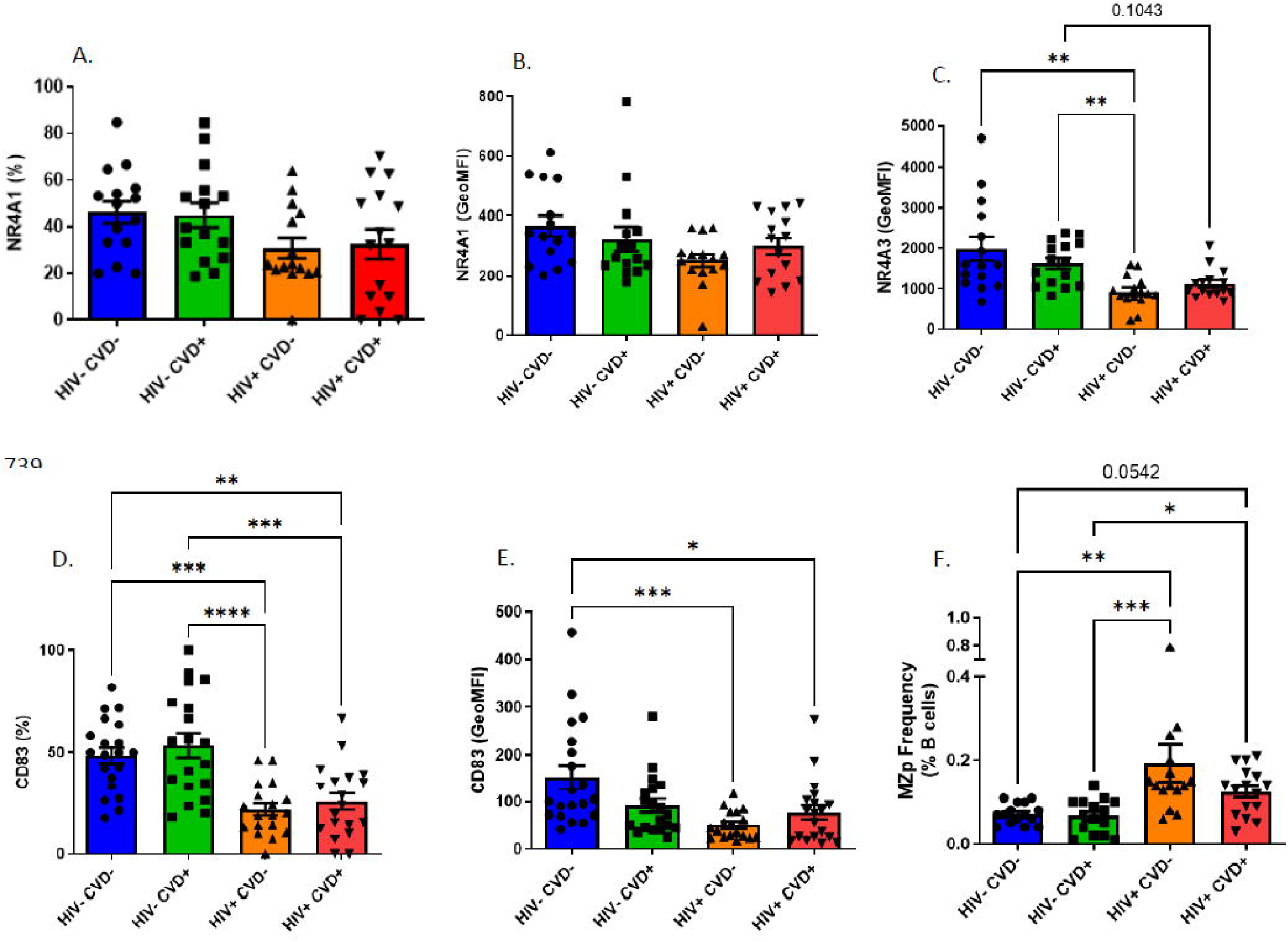
Expression levels of immunoregulatory markers NR4A1, NR4A3 and CD83 by blood marginal zone precursor-like (MZp) B-cells of HIV-uninfected (HIV-) and HIV-infected (HIV+) individuals of the CHACS. Expression levels of NR4A1, NR4A3 and CD83 by blood MZp of HIV uninfected participants, without and with cardiovascular disease (CVD) (HIV-CVD-, HIV-CVD+, respectively), and HIV-infected participants, without and with CVD (HIV+CVD-, HIV+CVD+, respectively) selected in sub-cohort (A). Shown are relative frequency of NR4A1 (A) and CD83 (D), as well as total relative frequencies of MZp in blood (F). MZp NR4A1 (B), NR4A3 (C) and CD83 (E) expression levels. Total relative frequencies of MZp in blood were assessed relatively to the percentage of total blood B-cells, and relative frequencies of MZp expressing NR4A1 and CD83 were assessed relatively to the percentage of total blood MZp B-cells. Expression levels were assessed with Geometric Mean of Fluorescence Intensity (GeoMFI). Normality was assessed with the Shapiro-Wilk test. A Kruskal Wallis test with a post-hoc Dunn’s test was used for testing statistical differences between groups in Fig. 3A, B, C, E and F. A One-Way ANOVA with a post-hoc Tukey was used for testing statistical differences between groups in Fig. 3D. * p < 0,05; ** p< 0,01; *** p< 0,001; **** p < 0,0001

Upon analysing blood MZp as to their expression levels of CD39 and CD73, we find that MZp from both HIV+ groups of sub-cohort (A) are the most affected, showing a downregulation of both ectonucleotidases (Figs. 4.A-D) independently of their CVD status. We also find that mature MZ from the blood of the HIV+ CVD+ group present a downregulation in CD39 and CD73 expression levels when compared to both the HIV-groups and to the HIV+ CVD-group (Figs 4.E-H).

**Figure 4:**
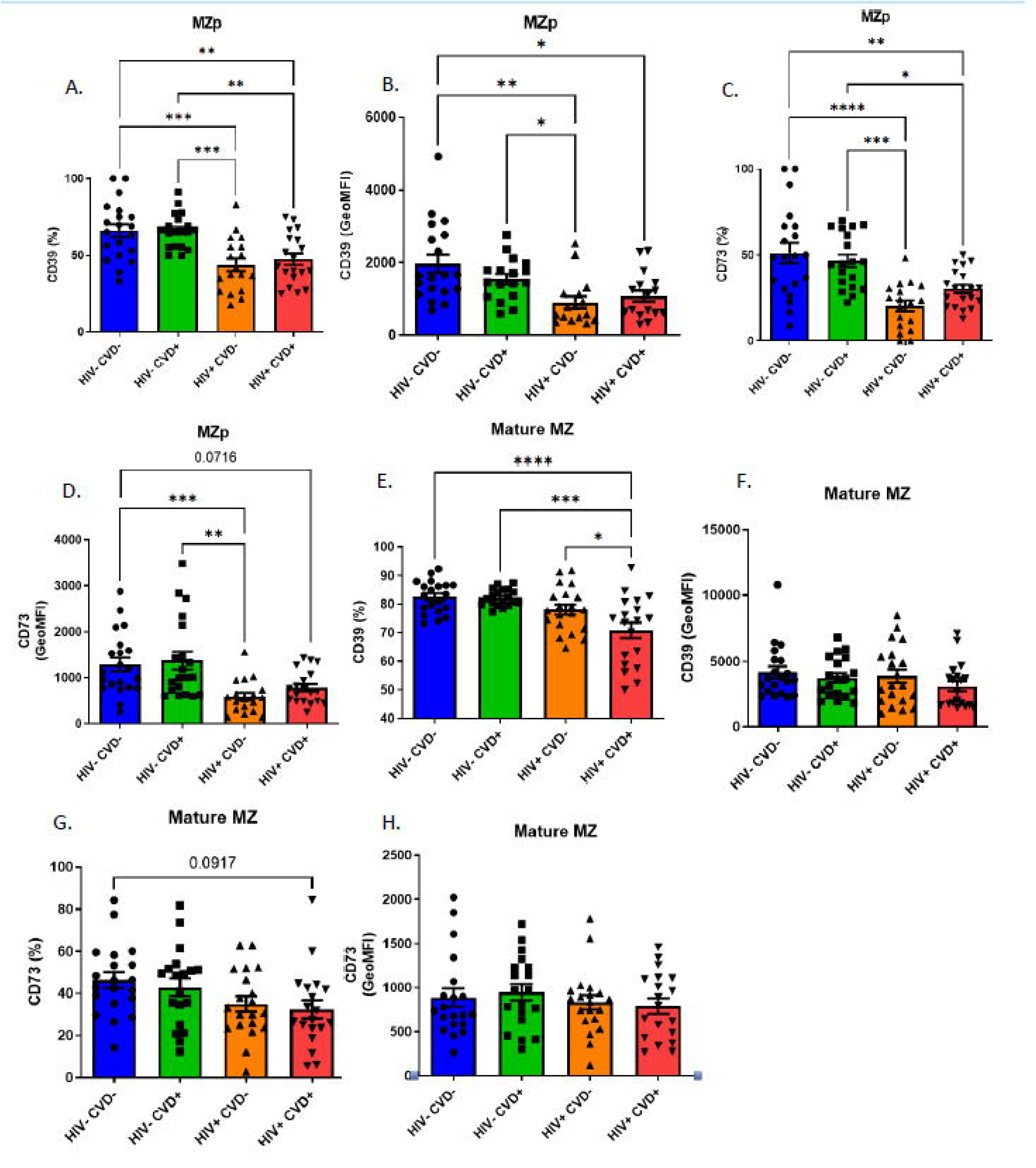
Immunoregulatory markers of the adenosine pathway, CD39 and CD73, are downregulated on blood marginal zone precursor-like (MZp) B-cells of HIV-infected (HIV+) individuals from the CHACS. Expression levels of CD39 and CD73 by blood MZp and mature MZ of HIV uninfected participants, without and with cardio-vascular disease (CVD) (HIV-CVD-, HIV-CVD+, respectively), and HIV-infected participants, without and with CVD (HIV+CVD-, HIV+CVD+, respectively) selected in sub-cohort (A). Shown are, relative frequencies of MZp and mature MZ expressing CD39 (A, E, respectively) and CD73 (C, G, respectively), as well as their CD39 (B, F) and CD73 (D, H) expression levels. Relative frequencies of MZp and mature MZ B-cells expressing CD39 and CD73 were assessed relatively to the percentage of total blood MZp and mature MZ B-cells, respectively. Expression levels for these markers were assessed with geometric mean fluorescence intensity (GeoMFI). Normality was assessed with the Shapiro-Wilk test. A One-Way ANOVA with a post-hoc Tukey was used for testing statistical differences between groups in Fig. 4A, C, E, G. A Kruskal-Wallis test with a post-hoc Dunn’s was used for testing statistical differences between groups in Fig. 4B, D, F, H. * p < 0,05; ** p< 0,01; *** p< 0,001; **** p < 0,0001.

As stated above, membrane CD83 expression levels are decreased in MZp from the blood of PLHIV from the CHACS, irrespective of their CVD status. As CD83 can also be found in a soluble form, which is normally involved in anti-inflammatory functions [34], we sought to investigate blood levels of soluble CD83 (sCD83) in individuals of the four study groups from sub-cohort (A). We find that the HIV-CVD+ group possesses the highest levels of sCD83 when compared to the other groups (Fig. 5.A). We also observe that levels of sCD83 correlate positively with soluble BAFF levels and with the BAFF/APRIL ratio in the HIV+ CVD+ group (Fig. 5.B and 5.C). sCD83 levels correlate negatively with APRIL levels in the HIV+ CVD+ group (Fig. 5.D), as with HDL levels (Fig. 5.E), while correlating positively with the LDL/HDL ratio in the HIV+ CVD+ group (Fig. 5.F). Altogether, our data show that in the context of long-term HIV infection, such as encountered in PLHIV from the CHACS, MZp Breg potential is still altered despite ART, suggesting that a sustained deregulation of MZp Breg potential and dampened immune surveillance may contribute to the premature development of atherosclerosis in these individuals.

**Figure 5:**
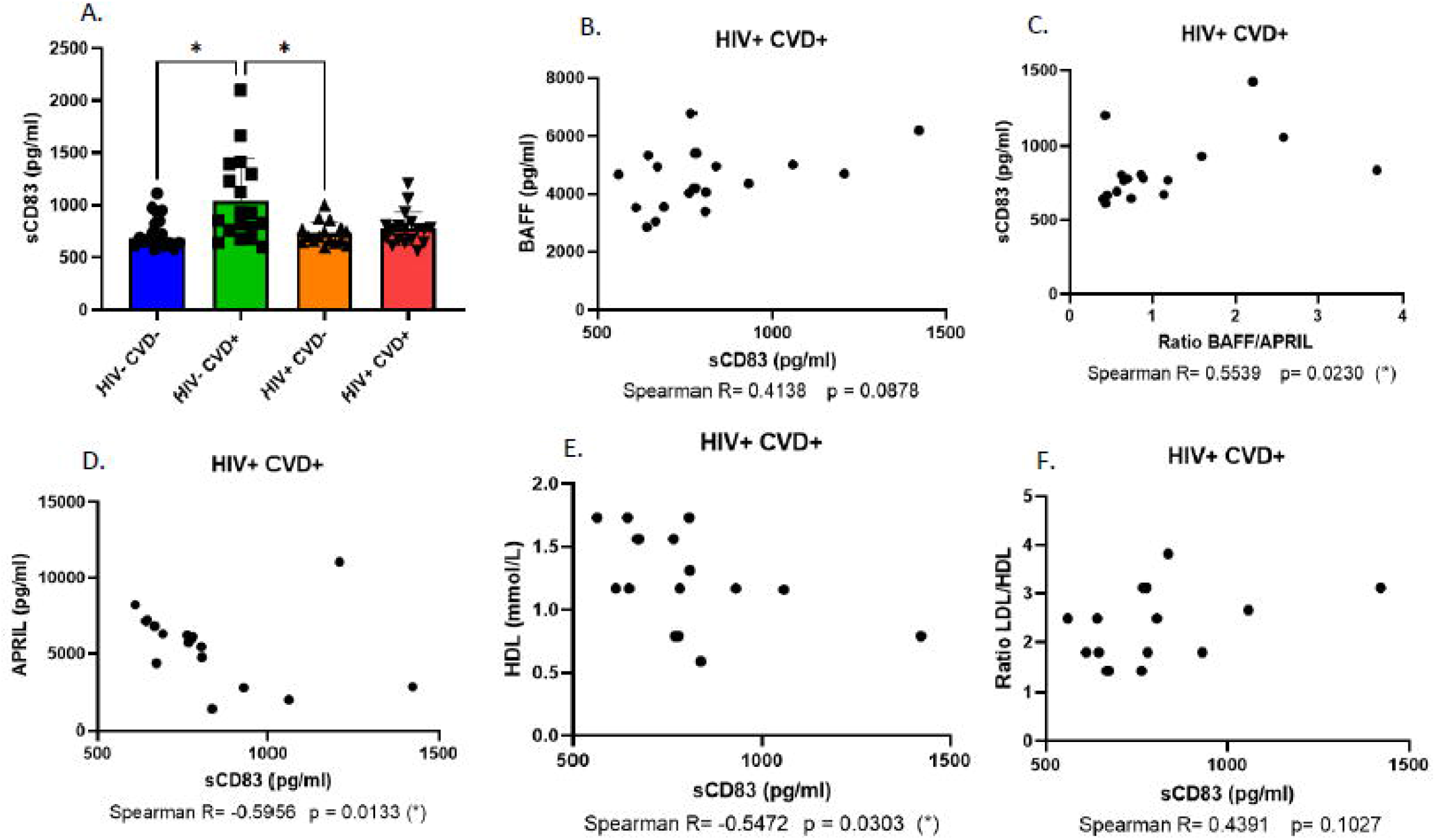
Soluble CD83 (sCD83) levels in the blood of individuals of the CHACS. Levels of sCD83 (A) in the blood of HIV uninfected participants, without and with CVD (HIV-CVD-, HIV-CVD+, respectively), and HIV-infected participants, without and with CVD (HIV+CVD-, HIV+CVD+, respectively) selected in sub-cohort (A). Correlation of sCD83 with soluble BAFF levels (B), BAFF/APRIL ratio (C) and APRIL levels (D). Correlation between sCD83 and HDL levels (E) and LDL/HDL ratio (F). Normality was assessed with the Shapiro-Wilk test. A Kruskal Wallis test with a post-hoc Dunn’s was used for testing statistical differences between groups in Fig. 5A. A Spearman correlation was used for testing for the correlations in Fig. 1B, C, D, E and F. CVD – Cardiovascular Diseases LDL – Low Density Lipoprotein; HDL – High Density Lipoprotein; * p < 0,05; ** p< 0,01; *** p< 0,001; **** p < 0,0001

### APRIL upregulates expression levels of Breg markers by blood MZp

Given the potential atheroprotective role of APRIL and its reported importance in modulating Breg profiles in humans [35-38], we sought to investigate whether APRIL could have an impact on the expression levels of Breg markers by blood MZp. To this end, human total B-cells from the blood of HIV-uninfected donors were cultured with different concentrations of soluble APRIL, and the expression of NR4A1, NR4A3 and IL-10 were analysed by flow-cytometry. We show that while 500 ng/ml APRIL only seems to slightly modulate NR4A1 expression levels by MZp (Figs. 6.A and 6.B), it significantly increases those of NR4A3 (Figs. 6.C and 6.D) and IL-10 (Figs. 6.E and 6.F). Interestingly, when soluble BAFF is added along with APRIL, this increase is diminished (Figs. 5.G and 5.I). Overall, our data suggests that APRIL can modulate the Breg potential of MZp, but this is impeded by excess BAFF.

**Figure 6:**
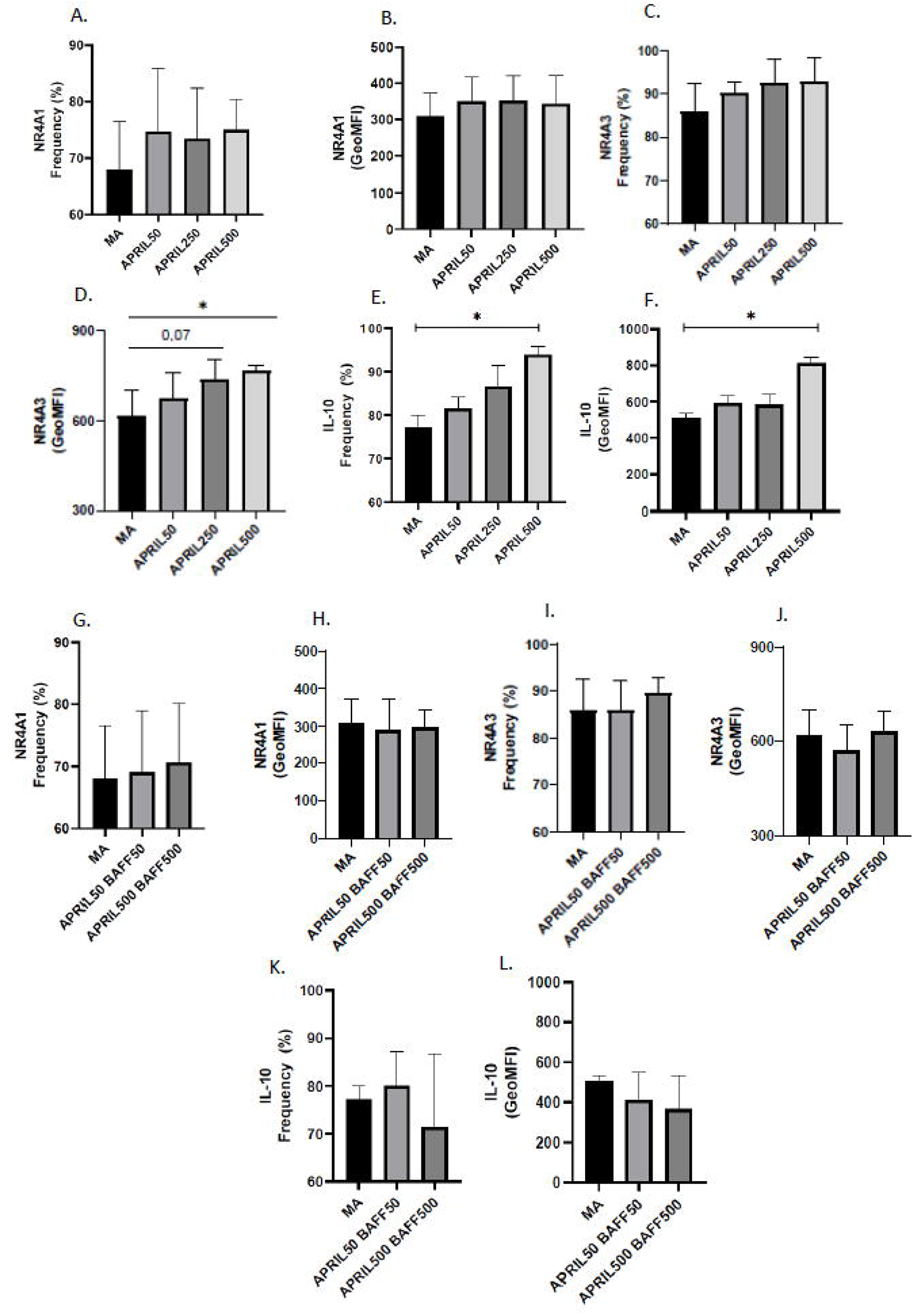
APRIL upregulates expression levels of immunoregulatory markers such as NR4A1, NR4A3 and IL-10 by blood marginal zone precursor-like (MZp) B-cells. Relative frequencies of MZp and their expression levels of NR4A1 (A, B), NR4A3 (C, D) and IL-10 (E, F) following treatment of total B-cells with APRIL at 50 ng/ml, 250 ng/ml and 500 ng/ml. Relative frequencies of MZp and their expression levels of NR4A1 (G, H), NR4A3 (I, J) and IL-10 (K, L) after treatment of total B-cells with BAFF and APRIL at 50 ng/ml and 500 ng/ml. Relative frequencies of MZp expressing NR4A1, NR4A3 and IL-10 were assessed relatively to the percentage of total MZp B-cells. Expression levels for these markers were assessed with geometric mean fluorescence intensity (GeoMFI). n= 3 HIV-uninfected donors. Normality was assessed with the Shapiro-Wilk test. A Kruskal-Wallis test with a post-hoc Dunn’s was used for testing statistical differences between groups in Fig. 6A-L. * p < 0,05; ** p< 0,01; *** p< 0,001; **** p < 0,0001

## Discussion

As stated earlier, HIV infection is marked by a chronic inflammation that persists despite ART [9-11]. Accordingly, we have shown that levels of BAFF are in excess in the blood of PLHIV from the Montreal PHI cohort, early on and despite 1-2 years ART. As such, we show in the current study that the excess of BAFF in the blood of PLHIV from the CHACS persists well beyond 15 years of ART, with levels of membrane-bound BAFF being more highly and significantly affected when comparing with soluble levels. This may be of importance and could reflect ongoing delivery of BAFF signaling in tissues recruiting these highly BAFF-expressing populations, either through its membranar or released soluble forms, as both were shown to deliver signaling [15]. Of note, in recent *in vitro* studies with tonsils from HIV-uninfected donors, we noticed that membrane-bound BAFF levels varied between each donor and this influenced MZp Breg capacities (Doyon-Laliberté, Aranguren *et al. unpublished observations*). This variation in membrane-bound BAFF was attributable to myeloid populations such as dendritic cells (DC), known to interact with B-cells in T-independent manners [14] and which could have thus modulated the tonsillar MZ B-cell environment. The fact that levels of soluble BAFF were still in excess in these individuals, despite long term ART, is reflective of a certain level of sustained systemic damage that may provoque the overall ongoing BAFF activity in the periphery. However, we noticed that excessive levels of soluble BAFF seem to persist on a smaller scale when compared to what we previously observed for other cohorts [13, 17]. This could be related to a certain degree of homeostasis reached after several years of controlled viremia. Indeed, several inflammatory markers associated with HIV infection such as IL-6 tend to normalise after long-term treatment [39], so it is also plausible that the same occurs with levels of soluble BAFF, to some extent. As such, we have shown that soluble BAFF levels fluctuate during the duration of the SIV infection in macaques [16]. Interestingly, HIV+CVD+ individuals are those who possess the highest levels of BAFF (in the two sub-cohorts studied in this work), suggesting a link between subclinical atherosclerosis and sustained excessive levels of this molecule. However, this hypothesis should be assessed in longitudinal studies.

The role of BAFF in atherosclerosis is complex and seems to differ between mice and humans. Evidence suggests that the neutralisation of BAFF is atherogenic in mice, whereas overexpression of BAFF in mice seems to be atheroprotective. The latter which appears to depend on TACI-expressing B-cells [40, 41]. Of note, MZ B-cells highly express TACI [14], supporting the atheroprotective role of this population. In humans, excess of BAFF has been found to correlate with CVD development in patients afflicted by autoimmune diseases such as systemic lupus erythematous (SLE) or Sjögren’s syndrome (SS) [42, 43]. Furthermore, BAFF has been found to be produced by adipocytes, and could be an adipokine linking obesity (one major cause of CVD development) with inflammation [44, 45]. Importantly, BAFF has been found to induce the apoptosis of precursor endothelial cells in the context of SLE, which is a sign of endothelial dysfunction, one of the key events implied in atherosclerosis development [46]. Despite being an analog of BAFF, APRIL is considered atheroprotective by its capacity to bind to HSPG such as perlecan, thereby blocking LDL retention and subsequent plaque formation [38]. Interestingly, APRIL has also been shown to favour the differentiation of Bregs, a role which could be involved in protection against atherosclerosis, as Bregs have also been shown to be atheroprotective in mice by dampening the local and systemic inflammation necessary to the development of the atherosclerosis plaques [35-37].

Our results thus suggest a possible link of excess BAFF in accelerating human atherosclerosis development in the context of HIV infection. Furthermore, our results provide some evidence for the possibility of the involvement of excess BAFF in the development of atherosclerosis even outside of the HIV context. Indeed, our data show that HIV-CVD+ individuals possess higher levels of membrane-bound BAFF than their healthy HIV-CVD-counterparts, which is consistent with the importance of inflammation in this context. In addition, our results demonstrate that BAFF levels correlate positively with many CVD risk factors, notably with TPV, while negatively correlating with CVD protective factors, in both HIV-CVD+ and HIV+CVD+ groups. Indeed, BAFF levels correlate positively with the Framinghan Risk Score in HIV-CVD+ but not in HIV+CVD+ individuals. The Framinghan Risk Score is the risk percentage of an individual of developing CVD in the next 10 years, based on CVD risk factors such as dyslipidemia, smoking, blood pressure, age and sex [47]. The fact that BAFF levels correlate positively with this score in HIV-CVD+ individuals indicates a link between BAFF levels and CVD development risk factors. Actually, the fact that BAFF is found to be augmented due to obesity and tabagism could be related to the fact that it correlates with the Framingham risk score[45, 48]. This does not seem to be the case for HIV+CVD+ individuals, where BAFF levels are found in greater excess due to the HIV context, and not necessarily due to CVD risk factors. However, this is in agreement with the fact that PLHIV have a higher risk of developing CVD when compared to HIV-uninfected individuals, even when matched for CVD risk factors [6].

APRIL has been shown to directly impede atherosclerosis formation [38]. In accordance with this atheroprotective role, we have shown that APRIL levels correlate negatively with CVD risk factors and positively with CVD protective factors both in HIV-CVD+ and HIV+CVD+ individuals. Interestingly, it was in the latter group that the strongest correlations were made. While we measured total APRIL blood levels by ELISA, Tsiantoulas *et al* demonstrated an atheroprotective role for the non-canonical APRIL, a form that preferentially binds to HSPG [38]. Further studies are required in order to measure this specific form of APRIL in the CHACS cohort, to assess its importance in HIV-related CVD development.

We had previously reported that APRIL was found in excess in the blood of PLHIV from the primary infection and up to one year of ART [13]. Differently from what we observed in past studies, APRIL has not been found in excess in HIV-infected individuals of the CHACS. The fact that APRIL returns to normal levels after years of treatment is in direct contrast with what has been found for BAFF levels, which are still elevated, especially in HIV+CVD+ individuals. This could represent an imbalance between BAFF’s atherogenic role and APRIL’s atheroprotective role. Accordingly, we report that the BAFF/APRIL ratio correlates positively with CVD risk factors while correlating negatively with CVD protection markers. This was true for both HIV- and HIV+ individuals. This not only further accentuates the importance of APRIL in protecting from CVD, but also indicates that BAFF atherogenicity outweighs the impact of APRIL’s atheroprotective role on CVD development in HIV-infected individuals.

Dyslipidemia is a common comorbidity found with PLHIV. As such, in untreated individuals, total levels of cholesterol, including HDL and LDL are downregulated. Following ART treatment, LDL and total cholesterol increases, while HDL is still downregulated [49]. The fact that high levels of BAFF seem to correlate with low HDL and high LDL could be related to the fact that these modulations are all related to the HIV context, and thus could reflect HIV disease progression. This could also explain APRIL’s positive correlation with HDL levels and negative correlation with the LDL/HDL ratio, as APRIL has also been found to be related to a slower HIV disease progression and a lower immune activation [50]. It is important to note that the CHACS does not control for statin usage. Nevertheless, as seen in Table 1, there are no difference in statin usage between our four study groups.

As mentioned earlier, the excess of BAFF in the HIV context correlates with deregulations in the B-cell compartment, notably with the MZp subset [13, 22]. Indeed, we have recently shown that blood MZp from HIV-infected individuals from the Montreal PHI cohort have a deregulated Breg profile and function, which is directly affected by excess BAFF (Doyon-Laliberté; Aranguren *et al. unpublished*). Considering that BAFF is still found in excess in HIV-infected individuals of the CHACS cohort, we expected to find deregulations in their MZp compartments. Given the role of Bregs in atherosclerosis surveillance [37], we also expected that these deregulations would differ between CVD- and CVD+ participants of the HIV-infected groups and HIV-uninfected groups. This hypothesis was found to be only partially true. Indeed, MZp from HIV+ individuals of the CHACS are still deregulated when compared to HIV-uninfected individuals even after 15 years of ART, as observed by the downregulation of NR4A3 and CD83 expression levels. Furthermore, CD39 and CD73 expression levels by blood MZp are also lower in HIV-infected individuals, similar to what we and others have reported for MZp and other B-cell populations [51, 52] (Doyon-Laliberté; Aranguren *et al. unpublished*). However, we have not found any differences in the expression levels for any of the Breg markers between the HIV-CVD- and HIV-CVD+ groups, despite BAFF being found to be relatively higher in the latter. Nonetheless, as stated above, this was not the case for the blood MZp from the HIV-infected groups. Also, in these groups, we noticed differences in CD39 and CD73 expression levels on mature MZ, where HIV+ CVD+ individuals expressed the lowest levels of these molecules, when compared to the HIV+ CVD-group. This could be of significance, given that the mature MZ are much more frequent in blood than MZp (Supplementary Figure 1). The adenosine pathway has been shown to be atheroprotective by dampening the inflammation needed to trigger atherosclerosis plaque formation [53, 54]. Thus, by having not only MZp, but also mature MZ expressing lower levels of CD39 and CD73, HIV+ CVD+ individuals are possibly burdened with higher inflammation levels than the other groups, likely contributing to early atherosclerosis development. Importantly, the loss of CD39 and CD73 has also been reported in regulatory T-cells of HIV+ CVD+ individuals of the CHACS cohort, further demonstrating the possible link between deregulations of the adenosine pathway and atherosclerosis [55].

Considering the downregulation of MZp membrane-bound CD83, we sought to measure sCD83 blood levels to assess its association with subclinical atherosclerosis. We observed that sCD83 levels are not reflective of MZp membrane CD83 expression levels. Indeed, there are no differences between HIV- and HIV+ individuals, and no correlation between sCD83 levels and MZp CD83 expression (data not shown). This is not surprising however, since CD83 is expressed by several cell populations, such as DCs (for which this molecule is an activation marker) and T-cells, whose level of CD83 expression might not have been affected by the HIV context [34]. We did, however, notice that HIV-CVD+ individuals possess higher levels of sCD83 in their blood, which could be in response to the inflammatory burden in these individuals. It is possible however that the greater inflammatory burden found in HIV-infected individuals precludes this attempt on regulation. As such, sCD83 correlates positively with BAFF and negatively with APRIL in the HIV+ CVD+ group, while also correlating positively with the BAFF/APRIL ratio and LDL/HDL ratio and negatively with HDL levels in this group. This demonstrates a possible pathological role of sCD83 in atherosclerosis development in HIV-infected individuals as well as a relation between sCD83 levels and HIV disease progression despite ART, as attested by the correlation between this molecule and dyslipdemia. Consistently, sCD83 has been found to be associated with certain pathologies such as malignancies and autoimmune diseases [56-59]. As such, more studies should be conducted to better understand the importance of sCD83 in human atherosclerosis and/or HIV infection.

APRIL has been shown to modulate human Bregs by allowing the generation of an IgA+ Breg population and by increasing IL-10 expression [35, 36]. Thus, we sought to investigate the role of APRIL in modulating the Breg potential of MZp, and found that APRIL increased NR4A3 and IL-10 expression by blood MZp *in vitro*. Further experiments will need to be conducted to assess if this increase translates to a stronger Breg function. Notably, when B-cells were co-cultured with APRIL and BAFF, the upregulation of Breg markers was dampened. We had previously reported that excess BAFF downregulates NR4As and IL-10 expression levels by MZp from HIV-uninfected donors (Doyon-Laliberté; Aranguren *et al. unpublished*). This suggests that in a context of relatively higher levels of BAFF than APRIL, the Breg potential of MZp is affected, impeding atherosclerosis immune surveillance.

## Conclusion

In conclusion, we have shown that PLHIV from the CHACS possess an excess level of BAFF in their blood even after 15 years of ART. We have also shown a positive association between BAFF levels and the presence of subclinical CVD, and a negative association between APRIL levels and subclinical CVD. Lastly, we have shown that MZp in PLHIV from the CHACS are still deregulated even after several years of treatment. Given the cross-sectional aspect of this study, we cannot determine a causal relationship between BAFF and APRIL and CVD development. However, our results suggest a link between levels of BAFF and subclinical coronary atherosclerosis in PLHIV from the CHACS, which need to be investigated in longitudinal studies. Strategies aimed at the modulation of BAFF and/or APRIL could thus be envisaged to dampen the inflammatory burden, restore Breg immune surveillance and prevent the premature development of CVD in PLHIV.

## Acknowledgements

We would like to thank Manel Sadouni, Marc Messier-Peet and Daniel Tremblay-Sher for the clinical data. We would like to thank Matthew Paniconi for the help in doing part of the correlations with BAFF and the clinical data. We would like to thank Mohammed Sylla and Olfa Debbeche for the preparation of the CHACS and healthy donor samples, respectively. We would also like to thank Dre Dominique Gauchat, Philippe St-Onge and Gaël Dulude from the CRCHUM flow cytometry platform. And lastly, we would like to thank all the patients from the CHACS cohort for their invaluable donation. The CHACS cohort is supported by a CIHR Team grant, Réseal FRQS-SIDA, Réseau de Bioimagerie du Québec and by the HIV Clinical Trial Network (CTN 272). MD receives a clinician-researcher salary award from the Fonds de Recherche du Québec – Santé.

## Conflict of interest

We disclose no relevant relationships.

## Notes

The authors have declared that no conflict of interest exists.

### Competing Interest Statement

The authors have declared no competing interest.

