## Supplemental Figures and Tables for "Subclinical Atherosclerosis is Associated with Discrepancies in BAFF and APRIL Levels and Altered Breg Potential of Precursor-like Marginal Zone B-Cells in HIV Treated Individuals"

**Supplemental Table 1 : Sociodemographic characteristics of the sub-cohort (B) used in this study**

| Group | HIV- CVD- (n=10) | HIV- CVD+ (n=20) | HIV+ CVD- (n=18) | HIV+ CVD+ (n=39) | p-value |
| --- | --- | --- | --- | --- | --- |
| Age | 52.93 (39.2 – 62.19) | 55.29 (42.79 – 70.57) | 51.77 (40.61 – 70.13) | 55.23 (44.22 – 74.43) | 0.28 |
| No. of male participants^1^ | NA | NA | NA | NA | NA |
| TPV (mm^3^) | 0 | 371.7 (34.0 – 1948.0) | 0 | 435.4 (6.74 – 1981) | 0.49 |
| LDL (mmol/L) | 3.54 (2.36 – 4.52) | 2.88 (1.67 – 5.44) | 2.99 (1.92 – 4.23) | 2.70 (1.15 – 4.57) | 0.051 (*)^2^ |
| HDL (mmol/L) | 1.33 (0.73 – 2.92) | 1.28 (0.89 – 1.99) | 2.99 (1.92 – 4.23) | 2.73 (1.15 – 4.57) | <0.0001 (****)^3^ |
| Median 10 years Framingham Risk Score (%) | 12.2 (2.00 – 18.00) | 11.89 (5.00 – 18.00) | 8.44 (4.00 – 18.00) | 10.23 (3.00 – 27.00) | 0.069 |
| Participants undergoing statin therapy (%) | 1 (10%) | 4 (25%) | 3 (17.64%) | 14 (35.9%) | 0.22 |

^1^All participants are male in this cohort

^2^Significant differences between HIV- CVD- and HIV+ CVD+ ( p = 0.04), as assessed by the Kruskal-Wallis test with post-hoc Dunn’s.

^3^ Significant differences between HIV- CVD- and HIV+ CVD- (p < 0.0001), HIV- CVD- vs HIV+ CVD+ (p = 0.0002), HIV- CVD+ and HIV+ CVD- (p < 0.0001) and HIV- CVD+ and HIV+ CVD+ (p < 0.0001), as assessed by the Kruskal-Wallis test with post-hoc Dunn’s.

**Supplemental Figure 1: Gating strategy for flow cytometry analyses.** After excluding doublets and dead cells, total B-cells (CD19+) were positively gated. Then, CD1c+ cells were selected, followed by IgM^+^ CD27^+^ double-positive cells. From the latter, the CD10- population was designated as mature marginal zone (MZ) B-cells, while the CD10+ population was designated as MZ precursors (MZp).

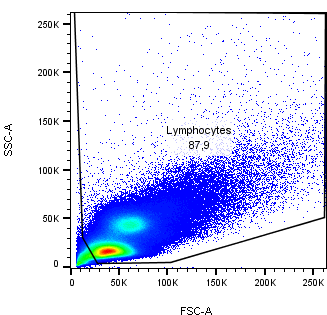

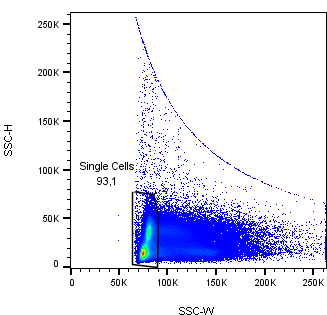

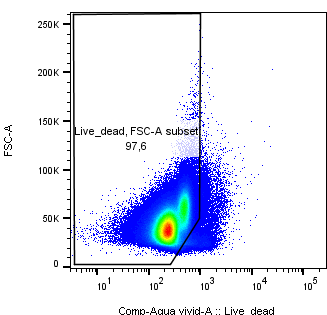

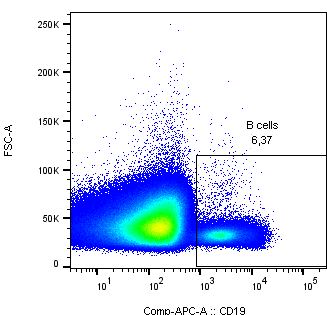

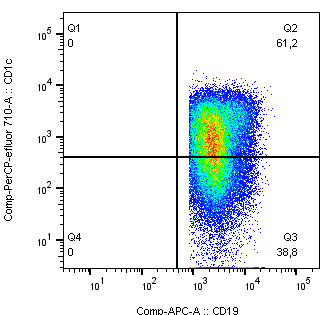

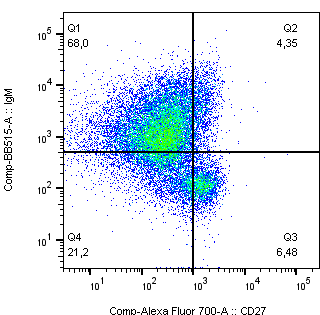

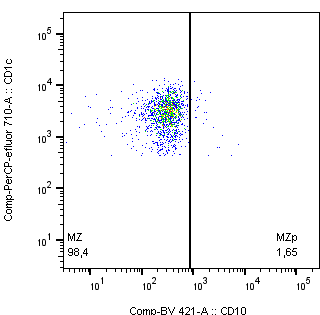

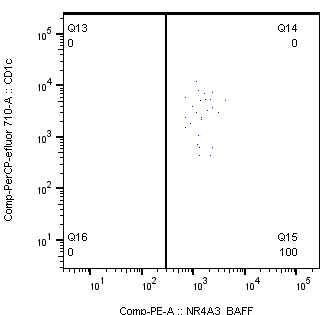

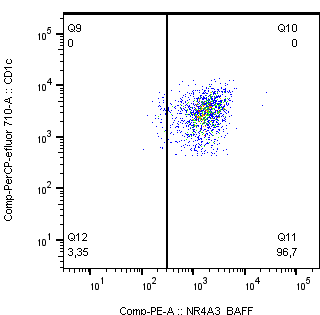

**MZp**

**Mature MZ**

**Supplemental Figure 2: BAFF levels and** **correlations between soluble BAFF and total plaque volume in the blood of HIV-uninfected and HIV-infected individuals of the CHACS sub-cohort (B).** Levels of soluble BAFF (A) in the blood of HIV uninfected participants, without and with CVD (HIV-CVD-, HIV-CVD+, respectively), and HIV-infected participants, without and with CVD (HIV+CVD-, HIV+CVD+, respectively) selected in sub-cohort (B). Correlation between soluble BAFF and total plaque volume in HIV+ CVD+ (B) and HIV- CVD+ (C) participants of the sub-cohort (B). Normality was assessed with the Shapiro-Wilk test. A Kruskal-Wallis test with a post-hoc Dunn’s was used for testing statistical differences between groups in Supp. Fig. 1A. A Spearman correlation was used for testing for the correlations in Supp. Fig. 2B, C. CVD – Cardiovascular Diseases. * p < 0,05; ** p< 0,01; *** p< 0,001; **** p < 0,0001

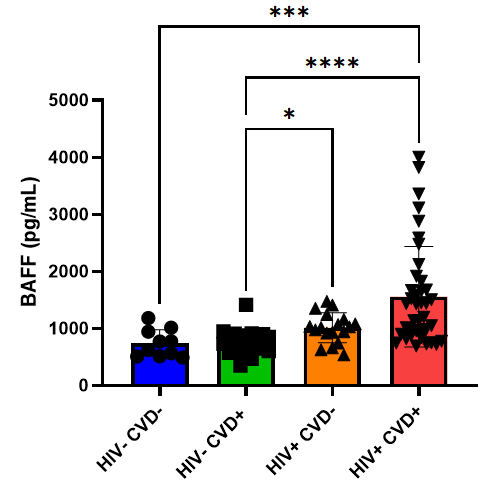

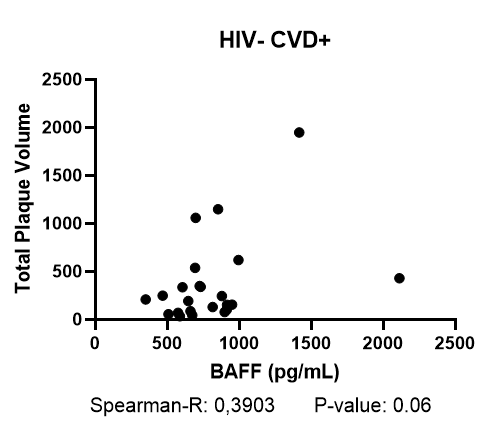

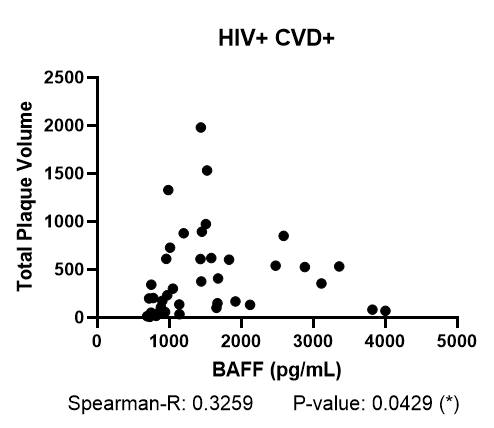

B.

C.

A.
